# DUET: a sequence-only ensemble for RNA–small molecule binding-site prediction

**DOI:** 10.64898/2026.10.28.762699

**Authors:** Kaifeng Xiong, Sulu Long, Chuanyu Lin, Jianhua Jia

## Abstract

Sequence-only predictors can score RNA–small-molecule binding sites without the need to obtain experimental RNA structures, but reported performance can be confounded by test-homologous training records and by inconsistent treatment of within-training redundancy. We introduce DUET, a sequence-only ensemble, and evaluate it with a test-clean protocol that controls these effects separately. Test-homologous records are removed, whereas sequence-identical training records are grouped into homology components and size-balanced to prevent family dominance. Across 10 paired seeds, DUET reached 0.240 ± 0.013 MCC, compared with 0.192 ± 0.031 for CoBRA-M3 trained on the same data (Δ = +0.047). Increasing the training pool from 80 to 290 records raised DUET by +0.135 MCC. DUET also reached 0.246 MCC on two independent SMARTPocket sets, supporting transport beyond the benchmark source.

**Graphical abstract:** The study separates benchmark overlap from model performance by removing test-homologous training chains while keeping the test set unchanged. DUET combines three sequence-derived predictors to assign nucleotide-level RNA–small-molecule binding probabilities directly from the RNA sequence.

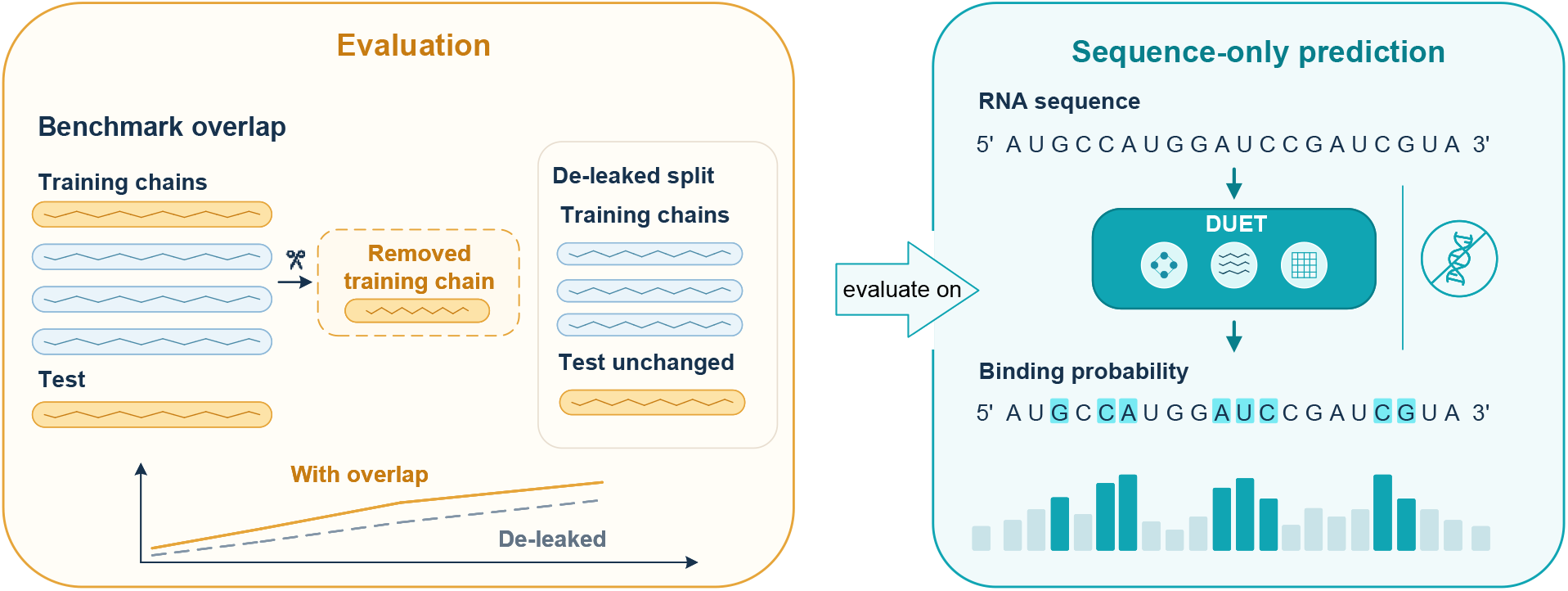

## 1 Introduction

Small molecules can target catalytic, regulatory and structural RNAs, extending drug discovery beyond protein-binding pockets (Kovachka et al., 2024; Jang and Shin, 2026; Nguyen and Le, 2026). A predictor of RNA–small-molecule binding sites assigns a score to each nucleotide so that candidate contact sites can be prioritized before experimental mapping. Structure-aware methods use three-dimensional geometry to describe pocket shape and chemistry (Oliver et al., 2020; Stefaniak and Bujnicki, 2021; Panei et al., 2024), whereas sequence-only methods use the nucleotide sequence and sequence-derived representations (Jang and Shin, 2026; Chen et al., 2022; Penić et al., 2025). The distinction matters operationally: structure-aware models can use information that is unavailable for most transcripts, while sequence-only models can be applied at transcriptome scale.

Benchmark construction in this field is not standardized. CoBRA removes sequence overlap with external tests before a random split (Jang and Shin, 2026), whereas RNAsite and HARIBOSS-derived structural splits use structural and sequence clustering to separate homologous RNAs (Su et al., 2021; Panei et al., 2022). These choices change which failure modes a reported score measures.

The evidence for comparing these two regimes is entangled with benchmark construction. Recent predictors draw training data from the Protein Data Bank through HARIBOSS (Panei et al., 2022), and the training resource is updated after benchmark publication. A test chain can therefore re-enter the training pool even when the benchmark was deduplicated when created. Such re-entry is a form of data leakage and a recognized threat to reproducibility (Kapoor and Narayanan, 2023). Removing all near-homologous training records is not a complete remedy because within-training redundancy carries additional supervision but can also dominate model fitting. Test-linked homology, within-training redundancy and the amount of retained training signal therefore need separate controls if model comparisons are to be attributed to model design.

We develop a test-clean, redundancy-aware protocol to isolate the prediction problem. Training or validation records are removed only when they reach at least 80 % identity with a canonical test chain. The remaining pool is partitioned into connected components at the same threshold, and whole components are isolated across cross-validation folds. Sequence-identical records are retained, and each record receives a sampling weight inversely proportional to its component size so that a large homology family cannot dominate training or model selection. Nested whole-component subsets further separate the effect of homology filtering from the effect of training-pool size.

We apply this protocol to DUET, a sequence-only ensemble of three predictors. A relational graph tower operates on frozen RNA-FM embeddings and edges derived from the sequence and its minimum-free-energy secondary structure. Two convolutional towers operate on frozen RiNALMo embeddings and mix local context with different inductive biases. No tower receives an experimental 3D structure, ligand identity, ligand coordinates or an experimental spatial neighbour graph, and the three output probabilities are averaged without joint training. DUET is therefore a sequence-only model with predicted secondary-structure context, not a 3D-structure-informed model.

The study separates three questions. First, how does DUET compare with CoBRA-M3 when both are retrained on the same test-clean pool, folds, thresholds and seeds? Second, how does the sequence-only model compare with GRASSP on shared internal benchmarks and on independent external sets? Third, how much of the observed performance is attributable to the retained training resource and to experimental three-dimensional topology? The comparisons are deliberately asymmetric in information access. GRASSP combines sequence-derived node features with a graph whose edges are constructed from a 3D RNA coordinate set, whereas DUET receives sequence-derived information only. The benchmark records are derived from experimentally determined RNA–ligand complexes, so the GRASSP graph describes a resolved RNA conformation rather than a feature that is normally available with a sequence query. We therefore use GRASSP as an information-rich reference and assess what sequence alone can recover, not as a like-for-like sequence-only baseline.

## 2 Materials and methods

### 2.1 Datasets, databases and resources

The task is ligand-agnostic, per-nucleotide binary classification. For each RNA chain, the model assigns a probability that a nucleotide belongs to a binding site for a small molecule. Ligand identity is not an input. We use the public benchmarks distributed with CoBRA (Jang and Shin, 2026). A nucleotide is positive when any of its heavy atoms lies within 4 Å of a ligand heavy atom. The released labels were used unchanged. TE18, TL12, JL10 and RB9 contain 17, 12, 10 and 5 chains after canonicalization, respectively, giving 44 canonical test chains. The raw training and validation partitions contain 345 and 43 chains. No experimental reagents, cells, organisms, or other biological materials were used. Table 1 summarizes the evaluation resource.

**Table 1.** Evaluation resource. Test-set counts are reported after removal of 5 exact cross-test duplicates. The test-clean pool is formed from the raw training and validation partitions and excludes only records reaching 80 % identity with a canonical test chain. Internal duplicates are deliberately retained for the sequence-redundancy-aware analysis.

| Partition | Source | Chains or records | Role |
|---|---|---|---|
| Raw training | HARIBOSS + TR60 | 345 | Pool input |
| Raw validation | HARIBOSS + TR60 | 43 | Pool input |
| Test-clean pool | HARIBOSS + TR60 | 290 | Training and grouped CV |
| TE18 | RNAsite (Su et al., 2021) | 17 | Canonical test |
| TL12 | ZeSTa (Gao et al., 2024) | 12 | Canonical test |
| JL10 | ZeSTa (Gao et al., 2024) | 10 | Canonical test |
| RB9 | RBind (Wang et al., 2018) | 5 | Canonical test |

The benchmark was designed to answer two separate questions: which records must be excluded to protect test independence, and how much redundancy should be retained inside the training pool. We treated only test-linked homology as leakage. Every raw training and validation record was aligned against every canonical test chain with global pairwise alignment. Pairwise identity was the number of matching positions divided by the number of alignment columns, and a pair was considered only when the ratio of sequence lengths was at least 0.8. A record was removed from the training pool when its maximum identity to any test chain reached 80 %. This removed 98 of 388 records: 86 from training and 12 from validation. We refer to the remaining 290 records as the test-clean pool.

Within the retained pool, sequence identity alone does not determine whether two records are interchangeable, because the same RNA sequence can occur in different ligand contexts. We therefore treated exact sequence matches as *sequence-identical records* and audited their labels rather than assuming that they provide identical supervision. The pool contains 180 unique sequences and 110 additional sequence-identical records, forming 50 groups. Only 5 groups (10 additional records) have identical labels throughout; 45 groups (100 additional records) contain label variation, and 85 additional records differ from the group’s modal label.

To prevent related records from crossing cross-validation folds, we built a sequence-identity graph over the pool at the same 80 % threshold and connected every pair whose identity reached the threshold. Labels did not influence component membership; they were used to audit whether sequence-identical records were interchangeable. After components were formed, their positive-residue counts were included when assigning whole components to folds. The graph had 115 connected components, including 61 singletons; the largest contained 27 records. Every component was assigned in full to one fold. We used 5 folds and balanced the number of validation chains to 58 per fold. The number of positive validation residues was not balanced because splitting a large homology component would violate the grouping constraint; all methods shared the same folds.

Within each training split, every record in a component of size *s* received sampling weight 1*/s*. Sampling was performed with replacement for as many draws as there were training records, so every component contributed equal total weight regardless of its duplication level. The weighting rule was fixed before fitting and applied equally to every model.

For each model, five-fold cross-validation used one common fold assignment. A model was trained on four folds, evaluated on the fifth at a fixed probability of 0.5, and the best epoch was recorded. The median of the five best epochs defined the final training length. Out-of-fold probabilities were pooled to select one decision threshold. The final model was then refitted from scratch on all 290 records for that epoch and evaluated at that threshold. This prevents the test set from entering epoch or threshold selection. Final comparisons used 10 paired training seeds.

The fixed-composition training-size analysis used nested whole-component subsets containing 80, 160 and 290 pool records. The test set, preprocessing and evaluation protocol were identical at every size. Three additional cluster-stratified fold assignments tested the stability of model selection, using five final seeds per assignment. A duplicate-weighting ablation compared grouped weights, uniform record weights and removal of sequence-identical records at fixed selected epochs and thresholds. Removing sequence-identical records also reduces the number of sampling draws per epoch. This ablation therefore combines record retention with a change in optimization exposure. These follow-up analyses are reported in the Supplementary Data.

## 2.2 DUET algorithm and software

DUET contains three independently trained towers whose output probabilities are averaged with equal weights (Figure 1). One tower reads a relational view; the other two read the same RiNALMo sequence representation with different local mixing patterns. All towers produce one logit per nucleotide, and no tower receives ligand identity, ligand atoms or experimental three-dimensional coordinates.

**Figure 1:**
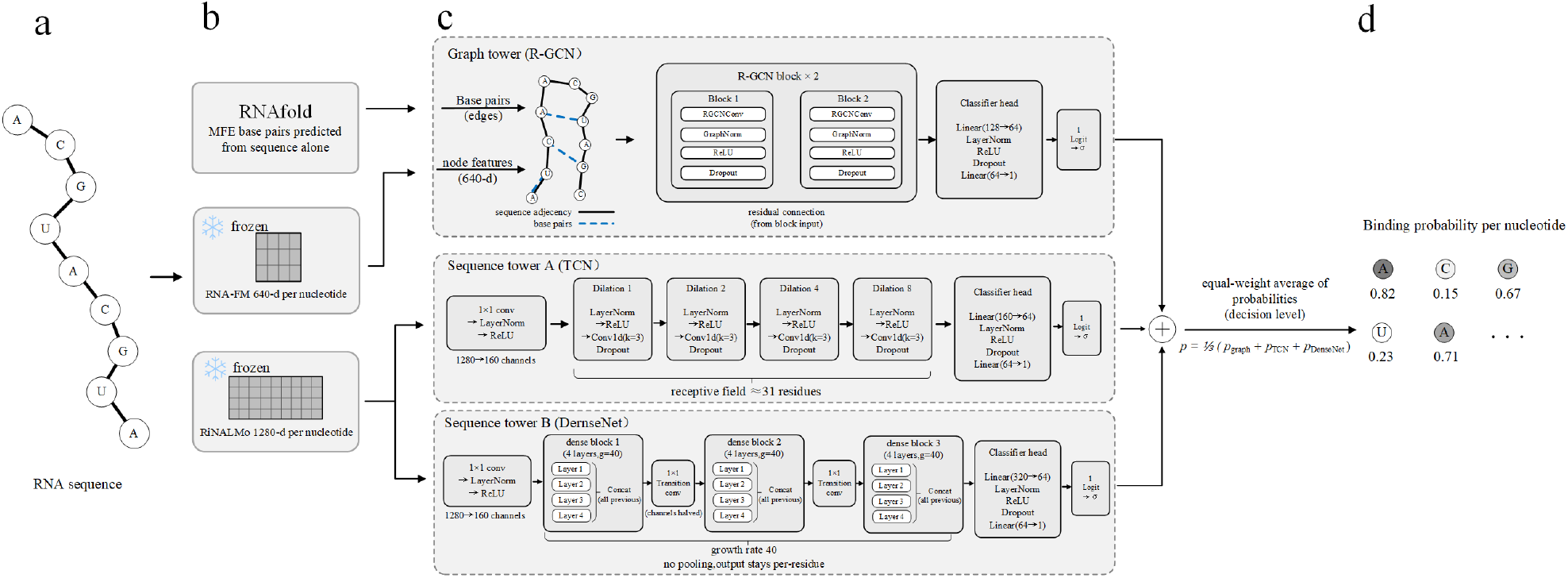
Architecture of DUET. (a) RNA enters as a nucleotide sequence. (b) RNAfold predicts a minimum-free-energy secondary structure from that sequence, and two pretrained RNA language models are used with their released weights fixed: RNA-FM for the graph tower and RiNALMo for the two sequence towers. (c) The graph tower propagates RNA-FM embeddings over sequence-adjacency and predicted-base-pair relations. The sequence towers use dilated temporal convolutions and dense convolutional blocks to mix local context while preserving per-nucleotide resolution. (d) The three binding probabilities are averaged at the decision level. The final output is a binding probability for each nucleotide.

The graph tower represents each chain as a multi-relational graph. Nodes are nucleotides with frozen RNA-FM embeddings (Chen et al., 2022); directed edges encode chain adjacency and RNAfold minimum-free-energy base pairs (Lorenz et al., 2011). Two relational graph convolution blocks (Schlichtkrull et al., 2018) use GraphNorm (Cai et al., 2021), residual connections and edge dropout (Rong et al., 2020). A small classifier maps the node representation to a binding logit. The graph tower encodes relational context that local sequence windows cannot capture.

Both sequence towers read frozen RiNALMo-giga embeddings (Penić et al., 2025). The temporal convolutional tower (Bai et al., 2018) applies a stem and four residual dilated blocks with increasing dilations, which gives each residue a local receptive field while keeping computation linear in chain length. The dense tower (Huang et al., 2017) applies a stem and three dense blocks with transition convolutions and no pooling, preserving nucleotide resolution throughout. The two towers use different inductive biases on the same embedding space and therefore provide complementary local views.

All three towers use the same combined objective. A class-balanced focal term (Lin et al., 2017) addresses the minority positive class, and a triplet-centre term (He et al., 2018) acts on the pre-classifier representation to compact each class and separate the two centres. The ordinary cross-entropy baseline and the complete training configuration are described in the Supplementary Data. The ensemble,

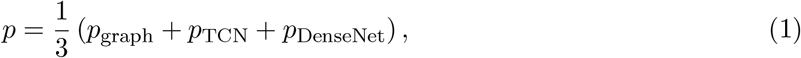

adds no parameters and requires no joint training.

### 2.3 Baselines and reference models

The primary baseline is CoBRA-M3, a released sequence-only model that maps RiNALMo embeddings to binding probabilities through a multilayer perceptron (Jang and Shin, 2026). We retrained it on the same pool, folds, sampling weights, epoch/threshold selection procedure and paired seeds as DUET. This controls for the benchmark update and makes the model comparison independent of released-training overlap.

GRASSP (Nguyen and Le, 2026) uses RNA language-model and predicted secondary-structure node features, with graph edges constructed from RNA 3D nucleotide coordinates. We reused its released graph assets for the internal benchmarks and reconstructed external graphs from deposited CIF coordinates after sequence alignment. We retrained two encoder pairings, ERNIE-RNA (Yin et al., 2025) and RNA-FM, using the authors’ public configuration fixed before our comparisons. The released graphs do not include an aligned RB9 set, so GRASSP was evaluated on TE18, TL12 and JL10, the three shared benchmarks containing 39 chains.

Matched R-GCN graph towers, shuffled-edge controls, the sequence+3D ensemble and the full GRASSP configuration grid are secondary analyses reported in the Supplementary Data.

### 2.4 External validation and statistical analysis

We used the SMARTPocket release (Thakare et al., 2026) to test transfer beyond the benchmark source. Chains reaching 80 % identity with the test-clean training pool or a canonical test chain were removed before evaluation. The independent sets contained 2 Jiang chains (97 residues) and 11 time-dependent chains (885 residues). The time-dependent release contains structures deposited after the HARIBOSS snapshot used for training. A HARIBOSS hold-out containing 6 chains (1010 residues) shares the training source and is reported only as a source-matched control. No external chain entered model selection, epoch selection or threshold selection.

MCC is the primary metric because binding nucleotides are the minority class in every benchmark. We also report AUC, AUPRC, F1, precision, recall and the geometric mean of sensitivity and specificity. Each training seed is evaluated at the threshold selected from grouped cross-validation. Method comparisons use same-seed paired differences and two-sided paired *t* tests. Holm correction is applied within prespecified comparison families: sequence-only comparisons on the four benchmarks, structure comparisons on the three shared benchmarks, crossed encoder/architecture controls, and independent external comparisons. We also report a paired chain bootstrap that keeps training seeds paired and resamples benchmark chains within each split, with 1,000 replicates. Chain resampling measures uncertainty across unseen RNAs; seed-level tests measure repeatability across training runs.

Absolute MCC is compared only between methods evaluated on the same chains. In particular, the label convention of SMARTPocket marks fewer positive residues than the four internal benchmarks, so its absolute values are not comparable with the internal scores.

### 2.5 Implementation and reproducibility

Pairwise sequence identities were computed with Biopython (Cock et al., 2009). Models were implemented in PyTorch 2.5.1 (Paszke et al., 2019) with PyTorch Geometric 2.6.1 (Fey and Lenssen, 2019). Metrics used scikit-learn (Pedregosa et al., 2011), and paired tests used SciPy (Virtanen et al., 2020). Training and inference were performed on a single NVIDIA RTX 3060 GPU. The repository contains the test-clean manifest, the complete identity graph, fold assignments, selected epochs and thresholds, trained checkpoints for all models and seeds, and external manifests. Scripts that regenerate the figures, main tables and Supplementary Data from the checked result files are included.

## 3 Results

### 3.1 Test-clean benchmark construction and matched sequence-only comparison

To separate test leakage from a change in available training information, we built a benchmark that protects test independence without discarding redundancy wholesale. Removing every near-duplicate could reduce both contamination and supervision, making model scores difficult to interpret. We therefore removed only training and validation records that reached 80 % identity with a canonical test chain. The canonical test set contained 44 chains after removal of 5 cross-benchmark duplicates, and 20 of these chains had at least one raw training or validation homolog. Removing 98 of 388 records left the 290-record test-clean pool (Figure 2a,b). The removed records comprised 86 training and 12 validation records; no label was changed.

**Figure 2:**
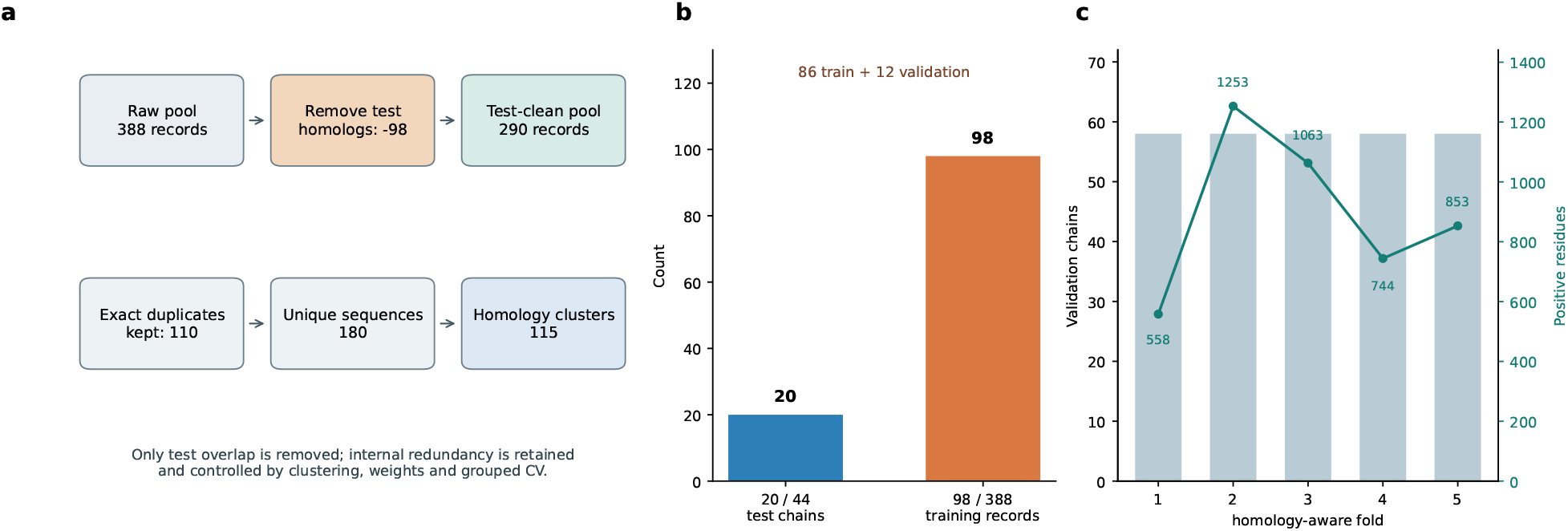
Test-clean construction and redundancy-aware cross-validation. (a) The raw training and validation partitions form a 388-record pool. Records reaching 80 % identity with a canonical test chain are removed, leaving 290 records. Sequence-identical records are retained and represented explicitly in the component structure. (b) 20 of 44 canonical test chains have at least one training or validation homolog. The 98 removed records comprise 86 training and 12 validation records. (c) The five folds contain equal numbers of validation chains but different numbers of positive residues; whole 80 % identity components are assigned to one fold.

Sequence-identical records were retained to keep de-leaking separate from de-duplication, not because they were treated as independent samples. The retained pool contained 110 additional sequence-identical records, representing 180 unique sequences, and 45 of its 50 sequence-identical groups contained label variation. This variation shows that an exact input match does not necessarily define the same ligand context. Retaining these records also changes the number of optimization draws, so we did not allow repeated sequences to act as uncontrolled weights. We used the 115 connected components of the 80 % identity graph to keep related records in the same validation fold and weighted each record by the inverse size of its component. Fold sizes were all 58 chains, while the number of positive validation residues varied from 558 to 1253 because whole homology components could not be split (Figure 2c). A shared fold assignment and component-balanced sampling gave every model the same data and prevented a large homologous family from dominating training.

In the primary matched comparison, DUET reached 0.240 *±* 0.013 MCC on the four benchmarks, compared with 0.192 *±* 0.031 for CoBRA-M3 (Table 2; Figure 3). The same-seed difference was +0.047 MCC (95 % CI +0.023 to +0.072), with Holm-adjusted *p* = 0.0073; DUET was higher in 10 of 10 seeds. The graph, TCN and DenseNet towers reached 0.193, 0.208 and 0.213 MCC. None of their individual differences from CoBRA-M3 survived the prespecified multiplicity correction, whereas the full ensemble exceeded the graph tower by +0.047 MCC (Holm *p* = 4.7 *×* 10^*-*4^).

**Table 2.** Sequence-only performance on the four benchmarks under the test-clean, redundancy-aware protocol. Values are means and standard deviations over 10 paired seeds. Each seed uses the epoch and threshold selected by grouped cross-validation.

| Method | MCC | Δ vs CoBRA-M3 | 95 % CI | Holm *p* |
|---|---|---|---|---|
| CoBRA-M3 | 0.192 ± 0.031 | — | — | — |
| DUET graph | 0.193 ± 0.017 | +0.001 | -0.023 to +0.025 | 0.9491 |
| DUET TCN | 0.208 ± 0.024 | +0.016 | -0.014 to +0.046 | 0.5391 |
| DUET DenseNet | 0.213 ± 0.012 | +0.021 | -0.005 to +0.047 | 0.3107 |
| **DUET ensemble** | **0.240 ± 0.013** | **+0.047** | +0.023 to +0.072 | 0.0073 |

**Figure 3:**
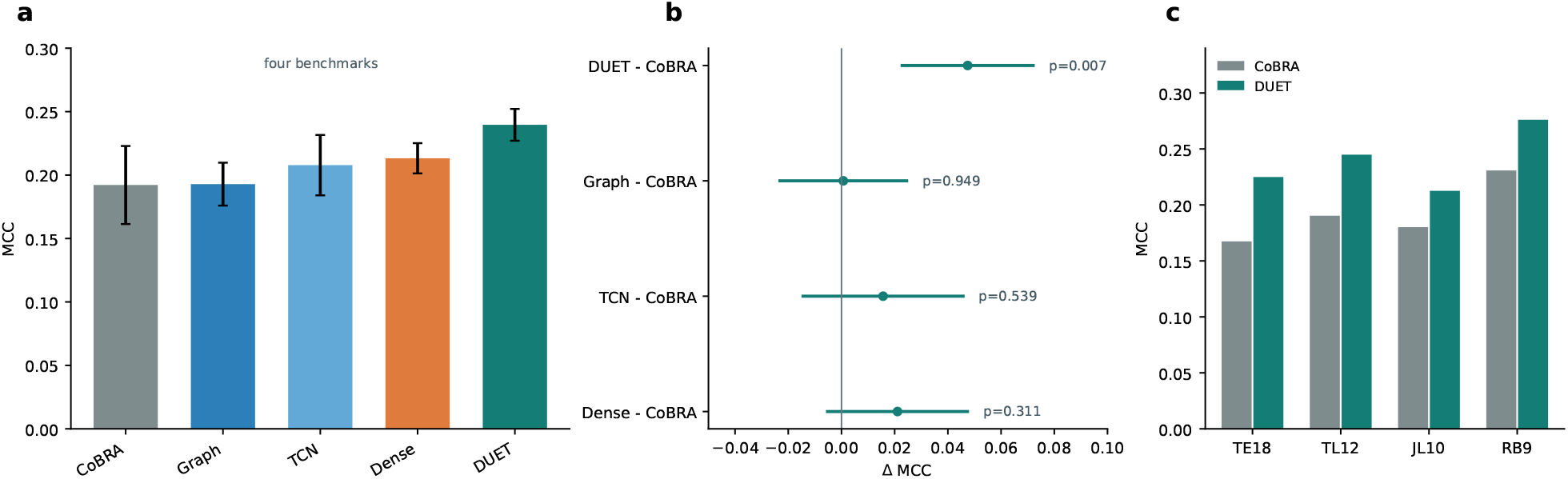
Internal model comparison and paired effects. (a) Mean MCC over 10 seeds for CoBRA-M3, the three DUET towers and the DUET ensemble on the four benchmarks. Error bars are one standard deviation over seeds. (b) Same-seed paired differences from CoBRA-M3 with 95 % confidence intervals. Labels give Holm-adjusted *p* values. (c) Per-benchmark MCC of CoBRA-M3 and the DUET ensemble.

For the same split, AUC increased by +0.027 (Holm *p* = 0.0280) and AUPRC by +0.035 (Holm *p* = 0.0146). F1, precision, recall and the geometric mean did not survive Holm correction.

A paired chain bootstrap kept the seed pairing and resampled benchmark chains. For the DUET-minus-CoBRA contrast, the 95 % interval was +0.020 to +0.077 MCC. The raw bootstrap *p* was 0.0020, and the family-wise bootstrap value was 0.0519. The seed-level conclusion is therefore more precise than the chain-level conclusion, consistent with the 44 test chains limiting the latter.

The direction also depended on the particular homology-aware partition. Across three additional cluster-stratified assignments, DUET minus CoBRA-M3 was +0.063 (Holm *p* = 0.1227), +0.077 (Holm *p* = 0.0236) and -0.008 (Holm *p* = 0.4199) MCC. We treat the canonical assignment as primary and use the additional assignments to estimate partition sensitivity, not as independent datasets.

### 3.2 Training-resource and redundancy sensitivity

To separate the effect of training-pool size from the identity filter, nested whole-component subsets were evaluated on the same test set. Performance increased monotonically from 80 to 290 records for every sequence model evaluated. DUET rose from 0.105 to 0.198 to 0.240 MCC, an increase of +0.135 from the smallest to the largest pool (Holm *p* = 1.7 *×* 10^*-*7^). CoBRA-M3 rose from 0.098 to 0.138 to 0.192 MCC (Figure 4a and Supplementary Data). The ordering between models cannot be inferred from the 80-record subset alone; the larger resource changes both absolute performance and the contrast between methods.

**Figure 4:**
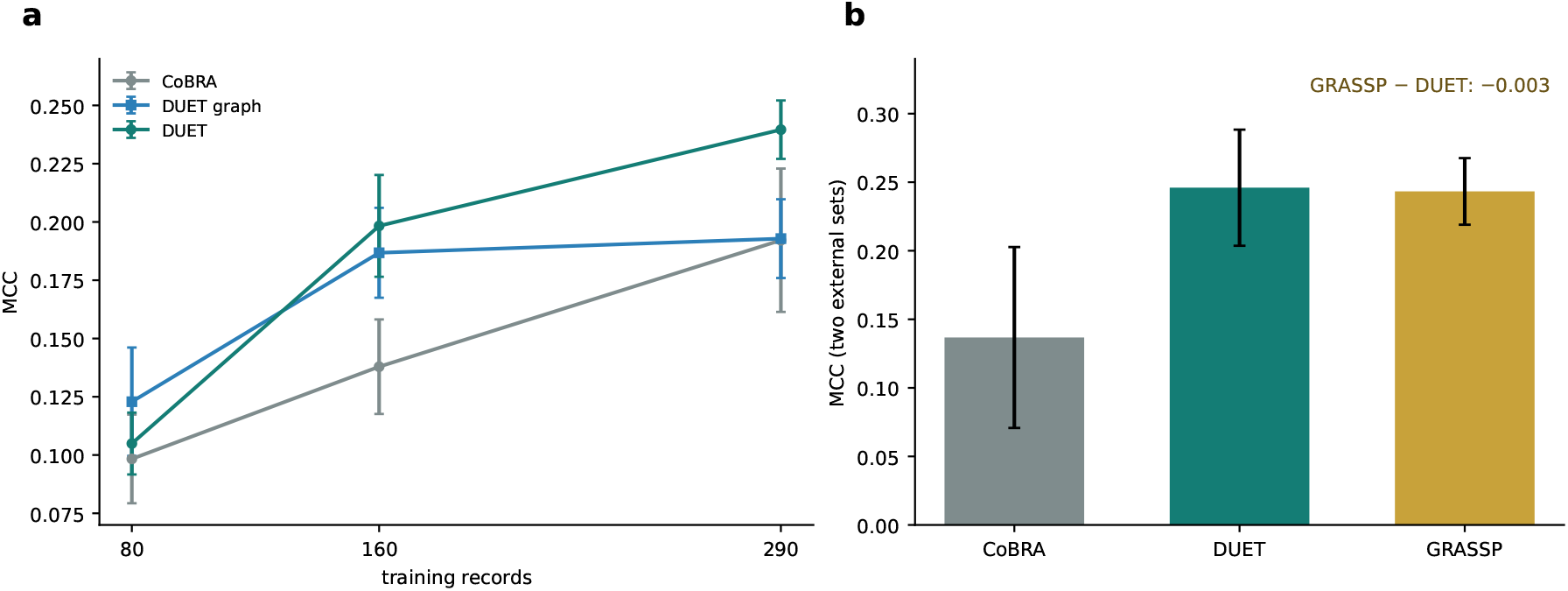
Training-size scaling and external transport. (a) Fixed-composition training-size curves on 80, 160 and 290 records with the same test set and evaluation protocol. Markers are means over ten seeds; error bars are one standard deviation. (b) MCC on the two independent SMARTPocket sets. GRASSP uses graph edges built from the deposited RNA 3D coordinates. These sets provide an external test of transport beyond the benchmark source.

The duplicate ablation used fixed selected epochs and thresholds. Removing sequence-identical records lowered MCC by -0.050 for the DUET graph tower (Holm *p* = 5.9 *×* 10^*-*4^). Switching from component-balanced to uniform record weights changed MCC by -0.011 (Holm *p* = 0.1979). Because removing records also reduces the number of sampling draws per epoch, the first contrast combines record retention with a change in optimization exposure. Results for GRASSP under the same ablation and the complete structural graph controls are reported in the Supplementary Information.

### 3.3 3D-informed reference and external transport

To contextualize sequence-only performance, we used GRASSP as a 3D-informed reference. GRASSP combines sequence-derived node features with graph edges built from a 3D RNA coordinate set, providing an information-rich reference for assessing what sequence alone can recover. On the three shared internal benchmarks, the authors’ default GRASSP RNA-FM model reached 0.261 *±* 0.014 MCC, compared with 0.227 *±* 0.016 for DUET. The seed-level difference was +0.033 MCC in favour of GRASSP (Holm *p* = 6.7 *×* 10^*-*5^), but a paired chain bootstrap gave a 95 % interval from -0.030 to +0.105 and *p* = 0.2857. The difference was consistent across initialization seeds, but its chain-level interval included zero.

The independent SMARTPocket sets test whether the ranking transfers beyond the benchmark source. DUET reached 0.246 *±* 0.042 MCC, GRASSP RNA-FM reached 0.243 *±* 0.024, and CoBRA-M3 reached 0.137 *±* 0.066. The GRASSP-minus-DUET difference was -0.003 MCC (95 % CI -0.038 to +0.032; Holm *p* = 0.9523), a difference whose confidence interval included zero. In the same comparison, DUET exceeded CoBRA-M3 by +0.109 MCC (Holm *p* = 0.0140). The source-matched HARIBOSS hold-out was excluded from the independent average. These external sets contain 2 and 11 chains, respectively, and provide a focused test of transport beyond the benchmark source. DUET reached comparable external performance to the 3D-informed GRASSP reference while retaining its matched-data advantage over CoBRA-M3.

## 4 Discussion

This study provides a sequence-only predictor and a controlled benchmark for evaluating it. Test-linked homologs can support memorization of a benchmark RNA, whereas repeated sequences inside the training pool can carry useful labels and close variants. The test-clean protocol removed 98 records with at least 80 % identity to a canonical test chain, retained 110 sequence-identical records, and grouped the remaining pool into 115 homology components. Under this shared construction, DUET exceeded same-data CoBRA-M3 by +0.047 MCC, with the same direction across 10 paired seeds. The result identifies a reproducible gain over a matched sequence-only baseline and supplies a reference point for future comparisons.

The training-resource experiment explains why benchmark construction changes model comparisons. Across nested pools from 80 to 290 records, DUET and CoBRA-M3 both improved, with DUET gaining +0.135 MCC. Absolute scores and model separations therefore depend on the amount of training signal retained after cleaning. At fixed epochs and thresholds, removing sequence-identical records lowered the graph tower by -0.050, whereas component reweighting had no significant effect. Record removal also reduced the number of optimization draws, so this contrast measures the retained resource rather than repetition alone. The prevalence of label variation among identical sequences argues against treating these records as simple copies, but it does not make them independent samples. We therefore use retention as the primary training-resource setting and report exact-sequence removal as a sensitivity analysis; the accompanying performance change cannot be interpreted as evidence that duplicates provide additional independent information.

The GRASSP comparison measures what sequence alone can recover relative to an information-rich 3D reference. GRASSP combines sequence-derived node features with predicted secondary-structure features and graph edges constructed from a supplied 3D RNA coordinate set; the benchmark graphs encode a resolved RNA conformation that is not available from sequence alone. On the three shared internal benchmarks, the authors’ default GRASSP RNA-FM configuration reached 0.261 MCC compared with 0.227 for DUET. The seed-level difference was +0.033 MCC in favour of GRASSP, while a paired chain bootstrap gave a 95 % interval from -0.030 to +0.105 and *p* = 0.2857. On the independent SMARTPocket sets, DUET reached 0.246 MCC and GRASSP RNA-FM reached 0.243 MCC; the 95 % interval for the GRASSP-minus-DUET difference was -0.038 to +0.032. These external sets contain 2 and 11 chains and provide a test of transport beyond the benchmark source. Sequence-only DUET therefore achieved comparable external performance without requiring 3D coordinates.

Matched graph controls and the full GRASSP configuration search are reported in the Supplementary Information. Those analyses quantify the added value of resolved geometry within this benchmark.

For prospective use, DUET is suited to first-pass residue-level scoring when only the RNA sequence is available or when the cost of structural preprocessing is prohibitive. Its primary operational advantage is input accessibility rather than reduced core-model cost. When a resolved 3D coordinate set is available, GRASSP provides an information-rich alternative; using it from sequence alone would require additional structure prediction and graph reconstruction, which was not evaluated here. Because DUET does not receive ligand identity, its output should be interpreted as a generic ligandability score for each nucleotide, not as compound-specific binding affinity or selectivity.

The benchmark itself is part of the contribution. Reporting only a final score hides the variables that determine the comparison. Studies of sequence-only and structure-aware RNA–small-molecule predictors should report test-linked homology removal, treatment of sequence-identical records, homology-component construction, cross-validation grouping, the final training-pool size and the availability of external structures. These quantities let readers separate a model effect from a change in the training resource. The same reporting framework can be reused as the underlying databases grow.

The evidence is bounded by the 44 internal test chains and the smaller external sets; the 80 % global-alignment criterion is an operational definition of sequence homology. Larger ligand-disjoint benchmarks and structure-matched comparisons will further improve precision and generality.

## 5 Conclusion

DUET provides a reproducible sequence-only baseline for predicting RNA–small-molecule binding sites. Under a test-clean, redundancy-aware protocol, it exceeded same-data CoBRA-M3 by +0.047 MCC. On the shared internal benchmarks, the paired comparison with GRASSP did not establish a performance difference (0.261 versus 0.227 MCC). On the independent external sets, DUET reached 0.246 versus 0.243 MCC, showing comparable performance without 3D coordinates. DUET therefore provides a practical sequence-only first-pass method, while GRASSP serves as an information-rich reference when resolved structures are available.

## Data Availability

The four benchmarks and the HARIBOSS+TR60 resource are distributed with the public CoBRA release (Jang and Shin, 2026). The SMARTPocket external sets are available from the public release (Thakare et al., 2026) at https://doi.org/10.64898/2026.06.18.732920. The canonical test set contains 44 chains after removal of 5 exact cross-test duplicates. The test-clean training resource contains 290 records, of which 180 have unique sequences and 110 are additional sequence-identical records. No raw label was changed. RNA-FM (Chen et al., 2022) and RiNALMo (Penić et al., 2025) were used with their released weights held fixed. Source code, processed graph tensors, manifests, identity tables, trained checkpoints and scripts are available at https://github.com/xiong-0212/DUET. The archived release is available at Zenodo with concept DOI 10.5281/zenodo.23635381.

## Supplementary data

Supplementary Data are available with this preprint. The supplementary file contains the leakage audit, complete metric tables, structure controls, training-size analysis, repeated grouped cross-validation, GRASSP sensitivity analysis, external validation and implementation details.

## Author Contributions

Conceptualization: J.J. Methodology: K.X. and J.J. Software: K.X. Validation: K.X., S.L. and C.L. Formal analysis: K.X., S.L. and C.L. Investigation: K.X., S.L. and C.L. Data curation: K.X. and S.L. Writing—original draft: K.X. Writing—review and editing: K.X., S.L., C.L. and J.J. Visualization: K.X. and C.L. Supervision: J.J. Project administration: J.J. All authors read and approved the final manuscript.

