## Supplementary Data for "DUET: a sequence-only ensemble for RNA–small molecule binding-site prediction"

#### 1 Scope and provenance

This supplement contains the complete numerical record behind the main manuscript. All values were generated from checked result files under `duet_clean_rebuild/results`. The main comparisons use 10 paired training seeds. Chain-level inference resamples benchmark chains and keeps seed pairing fixed. No result in this file uses a test chain for epoch or threshold selection.

The early tables are data-construction and leakage audits and do not involve model fitting. The remaining tables report models trained or refitted in this study under the documented test-clean protocols. External evaluations apply trained checkpoints without fitting on the external sets. No numerical result in these tables is transcribed from a prior publication.

#### 2 Relation to existing benchmark construction

Table S1 summarizes representative cleaning and split practices. The field does not use a single protocol: earlier structure-based benchmarks use structural and sequence clustering, whereas recent sequence-only studies often remove exact train–test overlap and then use an official or random split. Our protocol targets the latter setting while adding homology-disjoint cross-validation.

#### 3 Leakage and sequence-redundancy audit

Table S2 summarizes the construction of the test-clean resource.

The canonical test set contains 44 chains after removal of 5 exact cross-test duplicates: 17 in TE18, 12 in TL12, 10 in JL10 and 5 in RB9. 20 canonical test chains have at least one raw training or validation record at or above the identity threshold. All fold assignments used in the cross-model comparison share the same component partition. Table S3 reports the sequence-identical groups, and Table S4 lists the exact cross-test duplicates removed from the benchmarks.

#### 4 Internal performance and multi-metric inference

Table S5 gives MCC, AUC and AUPRC for all internal models. Table S6 gives the complete seven-metric paired comparison, including Holm-adjusted $p$ values and paired confidence intervals. These tables are generated directly from `results/followup/all_metric_stats/all_metrics.json` and the primary result JSON.

Table S1: Representative benchmark construction practices in RNA–small-molecule binding-site prediction. The summary follows the reported main protocols and is not an exhaustive survey.

| Study or resource | Test-linked redundancy | Cross-validation grouping | Within-training redundancy |
|---|---|---|---|
| RNAsite | $\text{TM-score}_{\text{RNA}} > 0.3$ structural clustering followed by sequence clusters | Train and test assigned by sequence cluster | Narrowed from 712 to 78 structurally distinct RNAs |
| HARIBOSS structural splits | TM-score structural clusters | Four structure-based train/validation/test splits | Similar structures assigned to the same split |
| CoBRA main split | Sequence overlap with external tests removed | Random 8:1:1 split; no homology-aware grouping reported | Exact duplicate policy not separated from split construction |
| GRASSP | Inherits benchmark-specific splits; overlapping sequences removed following CoBRA | Official benchmark-specific splits | Inherited from benchmark |
| MVRBind | Train60/Test18 and HARIBOSS structural splits | Inherited benchmark splits | Reported sequence/structure checks for selected external cases |
| **DUET** | Records at or above 80 % identity to any canonical test chain removed | Whole 80 % identity components assigned to one fold | Exact matches retained, audited and component-weighted; removal tested separately |

Table S2: Construction of the test-clean resource. The identity filter removes only records that reach 80 % global sequence identity with a canonical test chain. Exact sequence matches inside the retained pool are kept and represented by homology components.

| Quantity | Count |
|---|---|
| Raw training chains | 345 |
| Raw validation chains | 43 |
| Raw training + validation pool | 388 |
| Records removed for test homology | 98 |
| removed from training | 86 |
| removed from validation | 12 |
| Test-clean pool | 290 |
| Unique sequences in the pool | 180 |
| Additional sequence-identical records | 110 |
| Connected components at 80 % identity | 115 |
| Singleton components | 61 |
| Largest component | 27 |
| Folds | 5 |
| Validation chains per fold | 58 |

Table S3: Composition of the sequence-identical groups in the test-clean training pool. Group-level and record-level counts distinguish repeated supervision from sequence-identical records carrying different labels.

| Quantity | Count |
|---|---|
| Sequence-identical groups | 50 |
| Additional records in those groups | 110 |
| Groups with identical labels throughout | 5 |
| Additional records in all-label-identical groups | 10 |
| Groups containing label variation | 45 |
| Additional records in label-variable groups | 100 |
| Additional records differing from the modal label | 85 |

Table S4: Removal of exact sequence duplicates across the four test sets. A deterministic policy keeps the earlier benchmark in the order TE18, TL12, JL10, RB9 and then the lexicographically first chain identifier. 3 removed pairs had identical labels and 2 had different labels. Only the listed copies were removed from the test set; no training labels were changed.

| Removed chain | Source | Retained representative | Labels |
|---|---|---|---|
| `2tob_A` | RB9 | `2tob_A` (TE18) | identical |
| `4f8u_B` | TE18 | `2pwt_A` (TE18) | different |
| `1nem_A` | RB9 | `1nem_A` (TE18) | identical |
| `1q8n_A` | RB9 | `1q8n_A` (TE18) | identical |
| `2gdi_X` | RB9 | `7tzr_X` (JL10) | different |

Table S5: Internal performance from the ten-seed retraining under the large test-clean protocol. For each seed, each metric is first computed per benchmark and then averaged over the available benchmarks; MCC s.d. is computed across seeds. MCC, AUC and AUPRC are shown. GRASSP and its ERNIE-RNA crossed control are evaluated on the three shared benchmarks because the released RB9 assets are absent.

| Method | MCC | AUC | AUPRC |
|---|---|---|---|
| CoBRA-M3 (four sets) | 0.192 ± 0.031 | 0.657 | 0.451 |
| DUET graph (four sets) | 0.193 ± 0.017 | 0.641 | 0.450 |
| DUET TCN (four sets) | 0.208 ± 0.024 | 0.659 | 0.450 |
| DUET DenseNet (four sets) | 0.213 ± 0.012 | 0.666 | 0.454 |
| DUET ensemble (four sets) | 0.240 ± 0.013 | 0.684 | 0.485 |
| R-GCN seq+3D (three sets) | 0.212 ± 0.022 | 0.645 | 0.472 |
| R-GCN seq+ss+3D (three sets) | 0.180 ± 0.025 | 0.622 | 0.450 |
| GRASSP ERNIE-RNA (three sets) | 0.254 ± 0.018 | 0.671 | 0.494 |
| GRASSP RNA-FM (three sets) | 0.261 ± 0.014 | 0.686 | 0.509 |
| R-GCN ERNIE-RNA (three sets) | 0.161 ± 0.031 | 0.601 | 0.450 |

Table S6: All-metric paired comparisons and Holm-adjusted p values. Intervals are paired 95% confidence intervals over ten training seeds. The internal rows use the four benchmarks or the three shared benchmarks; external rows use the two independent SMARTPocket sets.

| Comparison | Metric | Delta | 95% CI | Holm p |
|---|---|---|---|---|
| DUET ensemble − CoBRA-M3 | MCC | +0.047 | +0.023 to +0.072 | 0.0073 |
| DUET ensemble − CoBRA-M3 | AUC | +0.027 | +0.009 to +0.044 | 0.0280 |
| DUET ensemble − CoBRA-M3 | AUPRC | +0.035 | +0.015 to +0.055 | 0.0146 |
| DUET ensemble − CoBRA-M3 | F1 | +0.050 | +0.004 to +0.096 | 0.1443 |
| DUET ensemble − CoBRA-M3 | Precision | +0.021 | -0.007 to +0.050 | 0.5144 |
| DUET ensemble − CoBRA-M3 | Recall | +0.059 | -0.042 to +0.159 | 0.6553 |
| DUET ensemble − CoBRA-M3 | GMean | +0.046 | +0.009 to +0.084 | 0.0822 |
| GRASSP RNA-FM − DUET ensemble | MCC | +0.033 | +0.025 to +0.042 | $6.7 \times 10^{-5}$ |
| GRASSP RNA-FM − DUET ensemble | AUC | +0.014 | +0.006 to +0.022 | 0.0105 |
| GRASSP RNA-FM − DUET ensemble | AUPRC | +0.015 | +0.003 to +0.028 | 0.0689 |
| GRASSP RNA-FM − DUET ensemble | F1 | +0.075 | +0.051 to +0.098 | $2.9 \times 10^{-4}$ |
| GRASSP RNA-FM − DUET ensemble | Precision | -0.035 | -0.050 to -0.021 | 0.0025 |
| GRASSP RNA-FM − DUET ensemble | Recall | +0.205 | +0.147 to +0.264 | $1.4 \times 10^{-4}$ |
| GRASSP RNA-FM − DUET ensemble | GMean | +0.049 | +0.032 to +0.066 | $6.7 \times 10^{-4}$ |
| External: DUET ensemble − CoBRA-M3 | MCC | +0.109 | +0.043 to +0.175 | 0.0140 |
| External: DUET ensemble − CoBRA-M3 | AUC | +0.073 | +0.031 to +0.115 | 0.0101 |
| External: DUET ensemble − CoBRA-M3 | AUPRC | +0.063 | +0.033 to +0.093 | 0.0033 |
| External: DUET ensemble − CoBRA-M3 | F1 | +0.080 | +0.036 to +0.125 | 0.0111 |
| External: DUET ensemble − CoBRA-M3 | Precision | +0.052 | +0.025 to +0.079 | 0.0067 |
| External: DUET ensemble − CoBRA-M3 | Recall | +0.135 | -0.035 to +0.305 | 0.3200 |
| External: DUET ensemble − CoBRA-M3 | GMean | +0.104 | +0.036 to +0.172 | 0.0214 |
| External: GRASSP RNA-FM − DUET ensemble | MCC | -0.003 | -0.038 to +0.032 | 0.9523 |
| External: GRASSP RNA-FM − DUET ensemble | AUC | +0.010 | -0.004 to +0.024 | 0.2560 |
| External: GRASSP RNA-FM − DUET ensemble | AUPRC | +0.039 | +0.010 to +0.069 | 0.0291 |
| External: GRASSP RNA-FM − DUET ensemble | F1 | -0.009 | -0.031 to +0.013 | 0.7597 |
| External: GRASSP RNA-FM − DUET ensemble | Precision | -0.023 | -0.046 to -0.001 | 0.0853 |
| External: GRASSP RNA-FM − DUET ensemble | Recall | +0.132 | +0.051 to +0.212 | 0.0198 |
| External: GRASSP RNA-FM − DUET ensemble | GMean | +0.017 | -0.005 to +0.040 | 0.2399 |

Table S7: Fixed-composition training-size curve. Nested whole-component subsets contain 80, 160 and 290 records with the same 44-chain test set and ten final seeds. Columns give MCC and the paired change from 80 records.

| Method | 80 | 160 | 290 | $\Delta_{160-80}$ | $\Delta_{290-80}$ |
|---|---|---|---|---|---|
| CoBRA-M3 | 0.098 | 0.138 | 0.192 | +0.040 | +0.094 |
| DUET graph | 0.123 | 0.187 | 0.193 | +0.064 | +0.070 |
| DUET ensemble | 0.105 | 0.198 | 0.240 | +0.093 | +0.135 |
| R-GCN seq+3D | 0.105 | 0.118 | 0.212 | +0.012 | +0.107 |
| R-GCN seq+3D ensemble | 0.111 | 0.198 | 0.239 | +0.087 | +0.128 |
| GRASSP RNA-FM | 0.134 | 0.220 | 0.261 | +0.086 | +0.127 |

Table S8: Independent SMARTPocket validation. The HARIBOSS source-matched control is excluded from the independent mean. Each value is the mean over ten paired seeds.

| Method | Jiang | Time-dependent | Independent mean |
|---|---|---|---|
| CoBRA-M3 | 0.183 | 0.091 | 0.137 ± 0.066 |
| DUET ensemble | 0.324 | 0.168 | 0.246 ± 0.042 |
| GRASSP RNA-FM | 0.328 | 0.159 | 0.243 ± 0.024 |

Table S9: Mechanism control for three-dimensional topology. All graph models use RNA-FM features and the same R-GCN. The shuffled graph preserves degree while rewiring endpoints. MCC is averaged over TE18, TL12 and JL10.

| Model | MCC |
|---|---|
| CoBRA-M3 | 0.179 ± 0.045 |
| DUET graph | 0.170 ± 0.020 |
| sequence + secondary structure | 0.180 ± 0.025 |
| shuffled 3D topology | 0.176 ± 0.026 |
| real 3D topology | 0.212 ± 0.022 |

Table S10: Repeated cluster-stratified cross-validation. Three alternative five-fold assignments were generated with the same 80% homology constraint. Five final seeds were used per assignment. The comparison is DUET ensemble minus CoBRA-M3 in MCC.

| Partition | Delta MCC | 95% CI | Holm p | Seeds better |
|---|---|---|---|---|
| 101 | +0.063 | -0.005 to +0.131 | 0.1227 | 4/5 |
| 102 | +0.077 | +0.034 to +0.121 | 0.0236 | 5/5 |
| 103 | -0.008 | -0.032 to +0.016 | 0.4199 | 2/5 |

Table S11: GRASSP hyperparameter sensitivity. All 24 author-grid configurations were screened on fold 0, and the top three were evaluated on the remaining folds. Final MCC and the paired difference refer to the three shared benchmarks; a negative difference means GRASSP is below DUET.

| Encoder | Selected configuration | OOF MCC | Final MCC | Delta | Holm p |
|---|---|---|---|---|---|
| ERNIE-RNA | lr=0.0001, batch=1, h=128, dropout=0.3 | 0.251 | 0.215 | -0.013 | 0.4185 |
| RNA-FM | lr=0.001, batch=16, h=128, dropout=0.3 | 0.243 | 0.208 | -0.019 | 0.0254 |

Table S12: Effect of sequence-identical record retention and component weighting. Grouped weights assign total weight one to each 80% homology component. Uniform weights assign one to each retained record. At fixed selected epochs and thresholds, the first contrast measures the combined effect of the record set and optimization exposure; the second isolates the weighting scheme.

| Model | Contrast | Delta MCC | Holm p |
|---|---|---|---|
| DUET graph | remove duplicates vs grouped weights | -0.050 | $5.9 \times 10^{-4}$ |
| DUET graph | uniform vs grouped weights | -0.011 | 0.1979 |
| GRASSP RNA-FM | remove duplicates vs grouped weights | -0.142 | 0.0105 |
| GRASSP RNA-FM | uniform vs grouped weights | -0.006 | 0.3868 |

### 5 Training-size and sequence-redundancy analyses

The fixed-composition curve uses nested whole-component subsets of 80, 160 and 290 records with the same test set and the same evaluation thresholds. It avoids the confound in a naïve identity threshold sweep, where changing the threshold also changes the amount of training signal. Component-balanced training assigns each component total weight one. The duplicate ablation fixes the selected epoch and threshold and varies the record set or component weights. Because removing records also reduces the number of sampling draws per epoch, this ablation measures a fixed-epoch resource change and does not isolate repeated supervision from optimization exposure. Table S7 and Table S12 contain the complete MCC contrasts.

### 6 Structural and robustness controls

The secondary graph towers used the RNA-FM backbone and classifier of the DUET graph tower. Sequence-adjacency edges were combined with experimental 3D neighbourhoods or with predicted base pairs. A degree-preserving shuffled graph retained the number of 3D edges while rewiring their endpoints. Each control used the same grouped cross-validation and paired seeds as the primary models.

The matched graph controls test whether experimental structure adds information beyond sequence and predicted secondary structure. The sequence+3D graph reached 0.212 MCC on the three shared benchmarks, compared with 0.180 for the graph without 3D edges, a difference of +0.042 (Holm $p = 4.5 \times 10^{-4}$). The degree-preserving shuffled graph reached 0.176 MCC. Real 3D edges exceeded this control by +0.037 MCC (Holm $p = 7.4 \times 10^{-5}$). Combining the sequence+3D graph with the two sequence towers reached 0.239 MCC, compared with 0.227 for sequence-only DUET and 0.261 for GRASSP on the same three benchmarks. These analyses are not used as inputs by DUET.

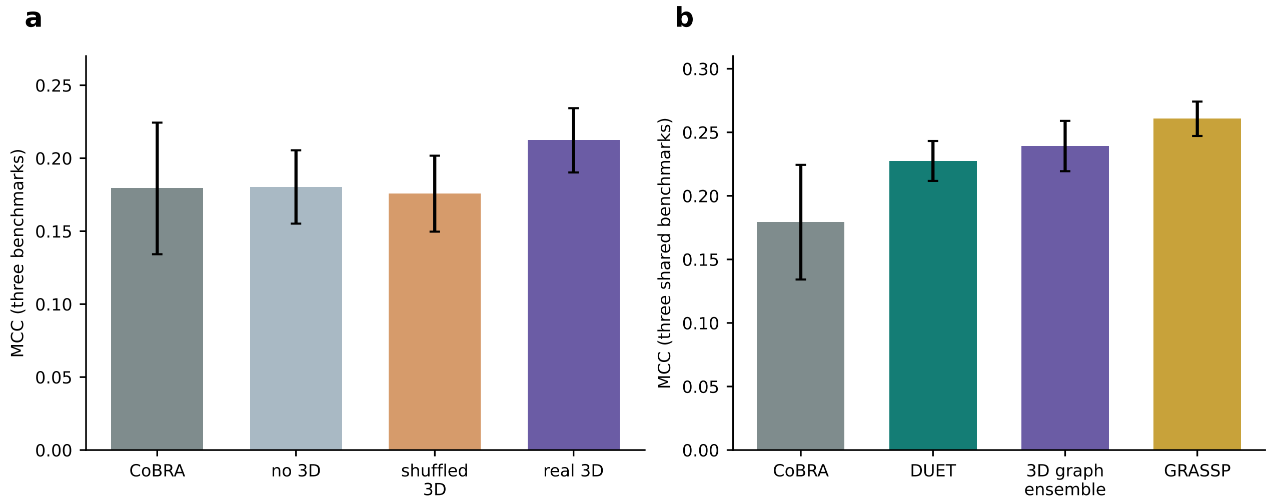


Figure S1: Structural controls on the three shared internal benchmarks. (a) CoBRA-M3, the graph without 3D edges, the degree-preserving shuffled 3D graph and the real sequence+3D graph. (b) CoBRA-M3, sequence-only DUET, the sequence+3D graph ensemble and GRASSP. Error bars are one standard deviation over 10 paired seeds.

Table S9 reports the degree-preserving shuffled-3D control. Table S10 reports three additional cluster-stratified five-fold assignments. These partitions are not independent datasets; they measure sensitivity to the particular homology-constrained fold construction. The default GRASSP RNA-FM comparison uses the authors' public training configuration, fixed before the shared-benchmark comparison and not tuned on those benchmarks. GRASSP node features combine RNA language-model embeddings with predicted secondary-structure features, whereas graph edges are constructed from RNA 3D nucleotide coordinates in structure files. The released benchmark graph assets were reused; external graphs were reconstructed from deposited CIF coordinates after sequence alignment. Predicted secondary structure is not a substitute for the 3D coordinate set. The sensitivity grid is reported separately in Table S11. All 24 author-grid configurations were screened on fold 0 and the best three were confirmed on the remaining folds. The grid is therefore a fixed-budget search, not a full five-fold evaluation of every configuration. The resulting GRASSP RNA-FM configuration reached 0.208 MCC on the shared benchmarks and the ERNIE-RNA configuration reached 0.215; the corresponding differences from DUET at 0.227 were -0.019 and -0.013.

### 7 External validation

Table S8 reports the two independent SMARTPocket sets and their mean. The Jiang set contributes 2 chains and 97 residues; the time-dependent set contributes 11 chains and 885 residues. The HARIBOSS hold-out (6 chains, 1010 residues) shares the training source and is excluded from the independent mean. External chains were excluded when they reached 80 % identity with the test-clean pool or a canonical test chain.

### 8 Selected epochs and thresholds

Table S13 reports the selected epoch and threshold for each model.

Table S13: Cross-validation selection. The final epoch is the median of the five fold-best epochs, and the threshold is selected from pooled out-of-fold predictions. The same selection rule is used for every model.

| Model | Final epoch | Threshold |
|---|---|---|
| CoBRA-M3 | 14 | 0.30 |
| DUET graph | 14 | 0.50 |
| DUET TCN | 18 | 0.50 |
| DUET DenseNet | 22 | 0.52 |
| DUET ensemble | 18 | 0.47 |
| R-GCN sequence+3D | 13 | 0.49 |
| R-GCN sequence+secondary+3D | 16 | 0.48 |
| R-GCN sequence+3D ensemble | 18 | 0.45 |
| R-GCN sequence+secondary+3D ensemble | 18 | 0.48 |
| GRASSP ERNIE-RNA | 81 | 0.50 |
| GRASSP RNA-FM | 79 | 0.49 |
| R-GCN ERNIE-RNA | 14 | 0.50 |

### 9 Downstream inference cost

We measured downstream model inference on 39 shared test chains with one CPU thread, using 5 timed repetitions after one warmup. Measurements start from precomputed frozen language-model features and graph tensors. They exclude language-model embedding extraction, RNAfold and GRASSP graph construction, so they quantify the trainable prediction network rather than the complete sequence-to-score pipeline.

| Model | Trainable parameters | ms per chain |
|---|---|---|
| DUET graph | 386,433 | 1.54 |
| DUET TCN | 524,897 | 2.34 |
| DUET DenseNet | 652,577 | 2.56 |
| DUET ensemble | 1,563,907 | 6.37 |
| GRASSP RNA-FM | 89,957 | 1.28 |

Table S14: Downstream inference cost on the shared test chains. The measurement excludes all frozen language-model feature extraction and all sequence or structure graph construction.

The three-tower DUET ensemble is larger and slower than the GRASSP prediction head in this downstream comparison. We therefore do not claim a computational-efficiency advantage. DUET's operational advantage is input accessibility, because it can be applied from sequence alone, whereas GRASSP requires a 3D RNA coordinate set before graph construction.

#### 10 Statistical definitions

For two methods evaluated on the same benchmark set and the same training seed, the paired effect is the mean within-seed difference across 10 seeds. The primary interval and test use the two-sided paired $t$ distribution. Holm correction is applied within each prespecified comparison family and metric. The chain bootstrap resamples chains within each benchmark, preserving the seed pairing and the cross-benchmark composition, with 1,000 replicates for the follow-up all-metric analysis. The bootstrap is reported as a secondary inferential route because the number of available benchmark chains is the limiting unit of evidence.

#### 11 Why a global low-identity split is not the primary analysis

Removing every training record above 70 % or 50 % identity also removes large amounts of non-test information. A model score then confounds two factors: reduction of residual leakage and reduction of the training distribution. We therefore use an explicit test-clean filter at 80 % identity, then estimate the effect of data quantity with nested fixed-composition subsets and estimate the effect of redundancy with the duplicate-weighting ablation. This design isolates the scientific question better than comparing models trained on progressively smaller pools with different class compositions. The complete manifests and identity tables are released so that alternative thresholds can be evaluated without ambiguity.

#### 12 Reproduction

The main numerical tables are generated by `scripts/41_export_paper_numbers.py`. The Bioinformatics figures are generated by `make_bioinformatics_figures.py` in this version. Raw results, selected epochs, thresholds and checkpoints are stored under `results/large_protocol` and `results/followup`. The commands and environment are documented in `HANDOFF.md` and `run_large_protocol.ps1`.
